# Science communication in online media: influence of press releases on coverage of genetics and CRISPR

**DOI:** 10.1101/2019.12.13.875278

**Authors:** Rafał Grochala

## Abstract

New scientific discoveries are communicated through multiple channels. Publications remain essential for scientists, whereas general public relies on news articles. Numerous studies investigated the path from publications to news and identified press releases as the key factor, associating them with issues such as exaggerations, but less is known about the direct influence of press releases on subject choice or on the content of media reports. Here, a cross-sectional sample of publications related to genetics and CRISPR is assessed in three independent datasets (n = 1362 publications, n = 461 press releases). Analysis finds 92.5% (CI = 88.5-96.5) dependence of news outlets on press releases in terms of topic choice and 39-43% explicit use of passages from press releases. Publications without press releases are described by 74x fewer news outlets. Even if they come from leading journals or universities, lack of a press release leads to 8.8x less coverage. Given the high impact of press releases, their reliability is especially relevant, but the majority of them omits interest disclosures - 84.3% (CI = 80.8-87.7) of press releases did not mention existing conflicts of interest, including multiple patent applications. These results establish the major indirect and direct role of press releases in science communication via online media. In line with previous research, this dependency raises concerns about possible distortions of science coverage.

## Introduction

Scientists intuitively view science communication as a one-way relationship in which they explain scientific findings to the knowledge-deficient public (Simis et al., 2016). This approach, known as the deficit model, was empirically rejected by studies in public understanding of science. Projects based on the deficit model aimed to increase public’s scientific literacy and interest, but did not reach their goals, did not influence policy or funding around science, as well as did not cope with climate change denial or anti-vaccine movements (Miller, 2001; Einsiedel, 2007; Peláez and Díaz, 2007; Kurath and Gisler, 2009; Short, 2013; Emery et al. 2014; Christiano and Neimand, 2017; Cook and Overpeck, 2018). Among those initiatives exercised in the 1980s and 1990s, science communication via media and through news was especially important. Frequent coverage of new scientific findings was expected to educate the public and elicit positive attitudes towards the science, but failed to achieve any measurable changes (The Royal Society, 1985; Miller, 2001). As a result, the science of science communication proposed another approach - public engagement with science - which was described as a move from “communication of settled science” to “communication about science, including its perils and pitfalls” (Jamieson et al., 2017). The role of scientific news in this model leans towards centers of engagement around which the public discusses new scientific findings, their details, credibility, and impact. Proliferation of internet aids the transition, enabling science communicators to directly engage with the public in social media, often in association with scientific news articles (Dudo, 2015; Kahle et al., 2016; Su et al., 2017; Hara et al., 2019; Jünger and Fähnrich, 2019).

News articles are relevant not only in modern science communication but also remain popular in the digitalized world. About 82% adults in the US regularly follow online news (Pew Research Center, 2019), with 36% of population reading multiple scientific news in each week (Pew Research Center, 2017). The majority of people rely on general news outlets on the internet, which are usually subdivisions of traditional news outlets. Therefore, difference between offline science journalism and online science journalism can be declared as trivial in principle (Su et al., 2015), with a few weighing-in factors: improved access to sources and experts, lack of some limitations (e.g. length, multimedia), and most importantly - higher pressure on journalists due to faster circulation of news and lower profitability of websites (Fahy and Nisbet, 2011).

Over decades, public coverage of science was condemned for uncritical approach to science (Williams and Gajevic, 2013) and science journalists were even named as retailers (Nelkin, 1995; Zhang, 2018) or priests (Murcott, 2009) of science. Media researchers attributed this stance to, among others, power imbalance between scientists and journalists (Fahy and Nisbet, 2011). The primary source of scientific news - peer-reviewed publication - is rarely comprehensible without sufficient expertise. Moreover, readability of publications decreases over time (Plavén-Sigray et al., 2017) and subjects of publications become more specialized (Casadevall and Fang, 2014). In consequence, journalists rely on dissemination of science by scientists themselves or by specialized teams at involved institutions. The latter evolved in the 1980s into public information offices (Rogers, 1986), which mediates between scientists and journalists. “Practitioners in the middle” make use of tools known in the public relations industry such as relationship development or distribution of ready-to-use materials (Göpfert, 2007). Among those, press releases are a critical tool, which provides a handy translation of a discovery from scientific to more general language.

Currently, 42% research institutes use press releases (PRs) and 64% scientists participate in press release preparation over a typical year (Entradas and Bauer, 2016; Marcinkowski et al., 2013). Journalists are known to widely rely on PRs to cover the science, with 75% - 84% news articles associated with PRs (de Semir, 1998; BBC Trust, 2011). Influence of PRs on news articles was investigated in terms of subject choice (Gonon et al., 2012), exaggerations (Sumner et al., 2014; Sumner et al., 2016; Bratton et al., 2019), quality (Kuriya et al., 2008; Woloshin et al., 2009; Brechman, 2011), language (Maat, 2007; Brechman et al., 2009; Wickman, 2013; Shea, 2015; Zhang, 2018), and industry funding (Woloshin and Schwartz, 2002). There were also anecdotal accounts of copying PRs by news outlets (e.g. Williams and Gajevic, 2013; Taylor et al., 2015). All work was based on top-down approach, where authors first collect news articles or PR. To date, literature did not assess this relationship in bottom-up approach, where publications are discovered first and this unbiased sample is assessed in association with PRs and news articles.

Here, traditional top-down (news to publications) approach is coupled with bottom-up (publications to news) method of discovery in three independent datasets with a total of 1362 publications promoted by 461 press releases and described by 11,862 news articles. The study measures the influence of press releases on science communication in terms of subject choice, direct use of PRs’ content, language, disclosures about competing interests, and explored in three popular cases. Subject choice was of particular interest, as it encapsulates essential question: who decides what scientific findings are communicated to the public through the news?

All analyzes are focused on news articles, press releases, and studies related to genetics or CRISPR. This topic was chosen to provide a cross-sectional sample, which encompasses basic science, technological advancements, applied science, as well as clinical trials. CRISPR specifically is also associated with high popularity in media and widespread conflicts of interests (Lim, 2018), presenting an interesting subject to study. At the same time, novelty of the field is associated with low presence in literature, which referred to coverage of CRISPR in a few narrative reviews (Gurev, 2017; Montoliu, 2018; Pei and Schmidt, 2019) and one recent analysis of 228 articles (Marcon et al., 2019). Present study takes into account over 865 peer-reviewed publications about CRISPR, which were described by 732 news articles.

## Results

To investigate communication of scientific discoveries in online media, a group of most popular news websites in the US was searched for articles related to genetics or CRISPR, published over twelve months. Between October 2018 and September 2019, there were 173 articles (12.4 per news outlet), which described 119 peer-reviewed publications. Thorough search revealed that 160 news articles (92.5%, CI = 88.5-96.5) and 110 publications (91.6%, CI = 86.5-96.7) were associated with press releases (PRs).

Two measures were employed to estimate the causal link between PRs and news articles. As a typical press release contains quotes (100/110 of collected PRs), their use in news articles provides direct evidence of press release’s influence on science communication. Manual assessment showed that 43.4% (CI = 35.8-51.0) of news articles used quotes from PRs. To assess the use of other parts of PRs, lexical similarity between full content of matched PRs and news articles was automatically measured with TAACO software and exceeded half of words in case of 39.3% (CI = 31.9-46.7) pairs. Almost all news articles in both groups were identical, suggesting that journalists often use other parts of PRs in addition to quotes. Estimate of 39-43% influence of PRs on news articles is conservative, as it did assess word-for-word similarity (not accounting for rewritten sentences), matched only public PRs (not accounting for private circulation of materials), and did not follow chains of sources (an article based on another article which in turn was based on press release). Those exceptions are partially investigated later in case accounts.

Similarity between headlines of PRs and news articles provided inconclusive results. Given limited space and narrow scientific dictionary, lexical similarity analysis and word2vector comparisons showed generally high overlap. However, there was no correlation between similarity of titles and similarity of content (Pearson’s r = 0.052, p = 0.52, Fig 1b), suggesting that headlines are often shaped by independent editorial decisions.

**Fig 1:**
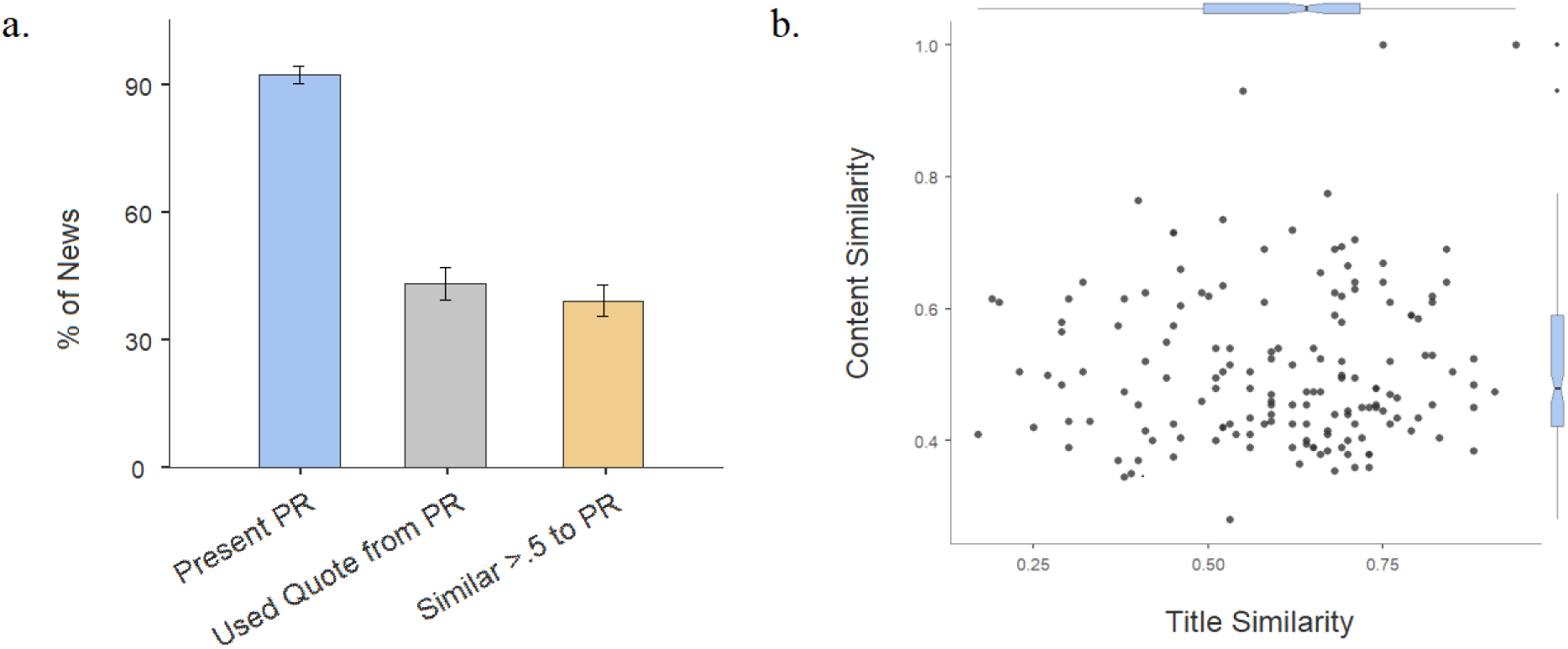
a. Fractions of news articles associated with press releases. b. Scatter plot of similarity between press release’s and news article’s title and content, which illustrates lack of correlation.

### High quality publications without press releases are not communicated

Initial exploratory analysis pointed to a high difference in popularity between press-released and not press-released publications: means of 60 and 25 news articles, means of 174 and 36 linking domains, means of 473 and 167 tweets. Even when publications were originally described by widely read news outlets (examples: BBC, New York Times), other online media rarely followed up in the absence of a press release. To check whether this observation could be attributed to newsworthiness (Rosen et al., 2016), popularity of news articles in both groups was compared. If not press-released publications were less popular due to lower newsworthiness, articles about them should also be less popular than articles about other publications. That was not the case in the comparison, which showed even slightly higher popularity of articles about not-promoted scientific publications (Supplementary Fig 1 and 2).

A new, independent dataset was used to precisely measure the effect of PRs on communication. In the same twelve-month period, 5 leading biomedical journals and 50 leading universities were searched for publications related to genetics or CRISPR. A total of 378 publications and 286 PRs was analyzed. Comparison of press-released and not press-released publications confirmed high difference in online media uptake, with an average of 8.8 more news outlets describing science promoted by PRs - 14.7 (CI = 12.5-16.9) news outlets in the press-released group in contrast to 1.67 (CI = 1.17-2.17) in not press-released (p<.00001, Fig 2). It is worth noting that publications overlooked by online media in the absence of PRs were from the most famous laboratories and universities in the field, including Feng Zhang laboratory (Scott and Zhang, 2017) or Robert Plomin works (Rimfeld et al., 2018), and were often published in widely-read, leading journals such as Nature or Science.

**Fig 2:**
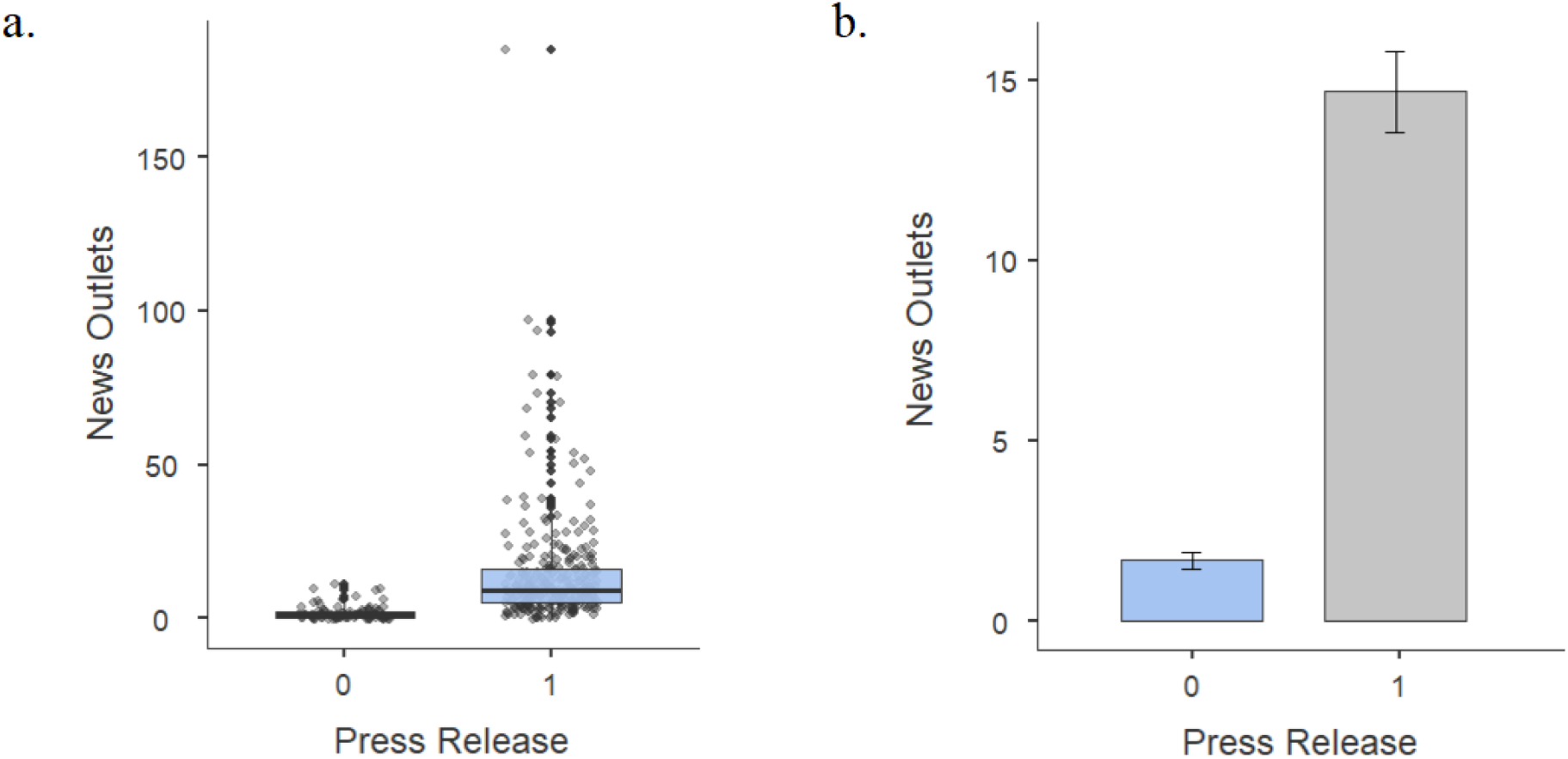
a. Box plots with all data points for 286 press-released and 92 not press-released publications from the high-quality dataset. b. Comparison of means for both groups.

To contrast this observation with an independent feature that should affect science communication, publications were grouped by their accessibility. In a comparison of 207 open access publications and 171 with paid access, there was no significant difference in news coverage, with mean of 12.1 news outlets (CI = 8.83-15.37) describing open access publications and 11.1 news outlets (CI = 9.22-12.88) covering publications without open access (p = 0.3, Supplementary Fig 3). Similar results were obtained after controlling for press release presence. This does not equal a lack of influence of accessibility on coverage, but it does prove at least lower influence. Given the sample size, effect size associated with the accessibility of publication is likely lower than d = 0.203 - whereas PRs have an effect size of d = 0.787.

### Replication of dependence on press releases in all CRISPR-related publications

To step outside of high-quality publications and focus on the whole field of CRISPR, another dataset of publications was collected, without a filter on journals or universities. Independent subset of 865 publications and 65 press releases was discovered in October and November 2019 - two subsequent months after the period of previous datasets. Consistently with previous results, press releases were associated with far higher media uptake: mean of 9.71 (CI = 6.79-12.63) news outlets in comparison to 0.13 (CI = 0.08-0.17) news outlets (p<.00001, Fig 3). Obviously, the primary function of a publication is to communicate results to other scientists, therefore there is no expectation that the public reads about most of the scientific findings. Moreover, 478/865 publications were virtually not present online, which is not described on any website and discussed in less than 5 tweets in social media. To test the influence of press releases acknowledging this property, not press-released publications were ordered by the highest popularity and 65 of them were compared to 65 press-released. Although the mean of news outlets in the not press-released group increased to 1.55 (CI = 1.13-1.97, similar to the high-quality group where CI spanned 1.17-2.17), it was still far different from press-released group with p<.00001. There was only one not-promoted publication, which reached the mean of 9.71 news outlets. Manual assessment showed that 57/65 most popular not press-released publications were described by specialistic blogs exclusively, instead of general online media.

**Fig 3:**
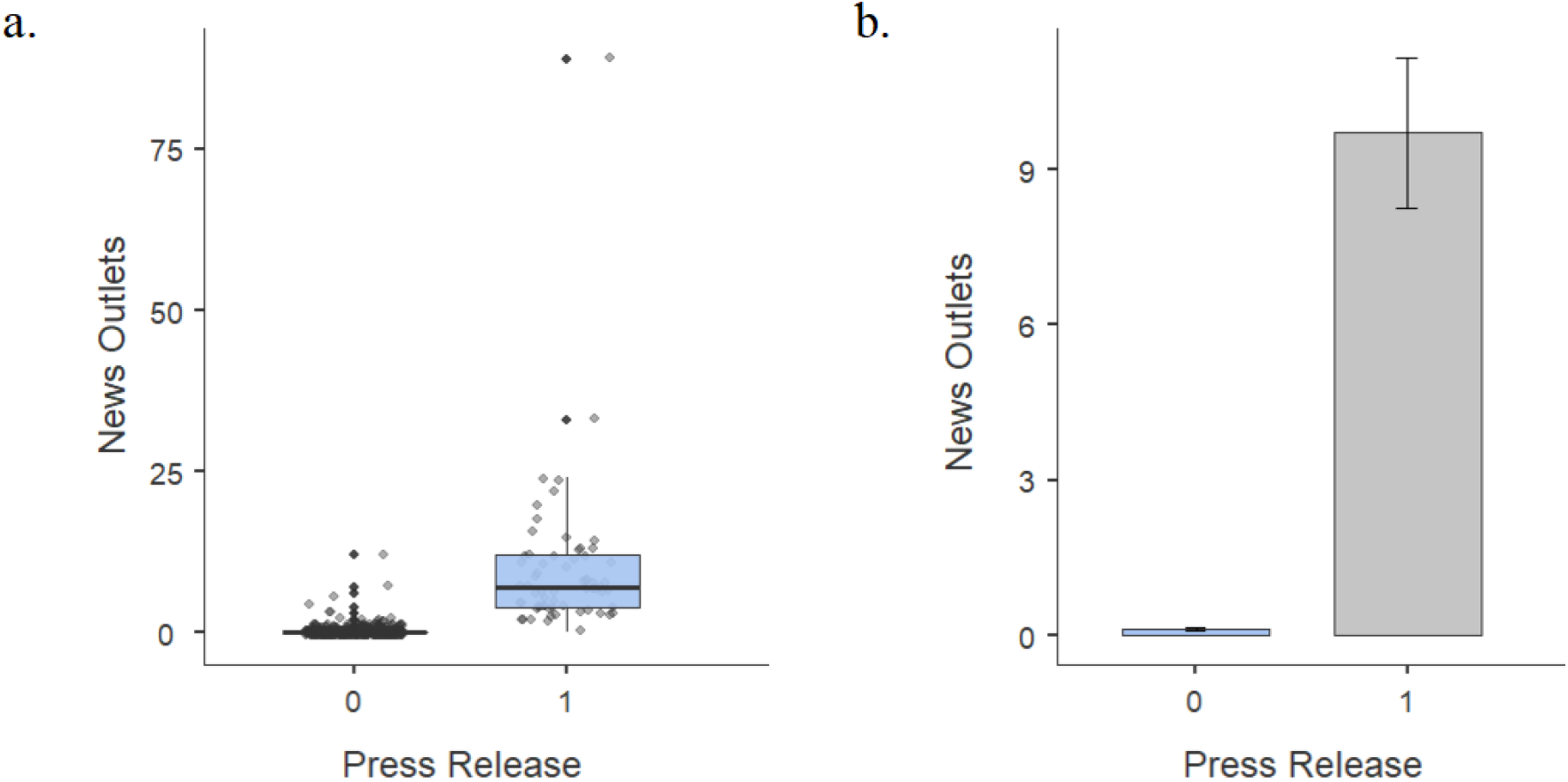
a. Box plots with all data points for 800 not press-released publications and 65 press-released publications from the third, unfiltered dataset. b. Comparison of means in both groups.

### Conflicts of interests are rarely communicated in press releases

Demonstrated importance of PRs in science communication prompted further investigation of PRs’ features. As CRISPR-related publications are known for a multitude of competing interests, including patent applications (Lim, 2018), it would be interesting to verify how it is acknowledged in PRs. In the dataset of high-quality publications (leading journals and leading universities), 203 publications explicitly declared no conflicts of interest and 150 listed competing interests. Out of those 150 publications, 127 were associated with PRs, but conflicts were communicated to the public only in 20/127 cases - 84.3% (CI = 80.8-87.7) did not communicate existing conflicts of interest. It is worth noting that the remaining publications were mostly not open-access, allowing journalists to freely and legally discover competing interests only in 30 more publications.

As conflicts of interests ranged from minimal (such as repaid travel costs) to significant (such as patents), publications were further divided by the type of competing interests. There were 71 publications that listed patents closely related to described work - often, methods and results of the publications were themselves the subject of a patent application. Of those, 58 publications were promoted by PRs, which failed to mention the patenting process in 45 cases (77.6%, CI = 66.6-88.6).

To explore the possible reason for omitting such important information, a hypothesis was tested, where disclosure of competing interests could discourage journalists, leading to lower media uptake. Although limited by sample size - because PRs mentioning conflicts were rare - initial comparison did not find significant difference in media coverage, with means of 17.0 (CI = 12.45-21.55) news outlets for avoided disclosures and 20.2 (CI = 10.66-29.74) for written disclosures (p = 0.53, likely detection of minimum effect size d = 0.481, Supplementary Fig 4).

### Language of press releases

As PRs are associated with the majority of science communication, their use of words could influence online media uptake. Two measures of wording known from the literature were employed to evaluate PRs: the intensity of scientific jargon (Rakedzon et al., 2017) and presence of promotional keywords (Maat, 2007). There was low correlation between jargon scores and popularity among online media (Pearson’s r = 0.168, p<.05, Supplementary Fig 5) and no significant correlation between promotional words and popularity (Pearson’s r = 0.085, p = 0.15, Supplementary Fig 6). It is not synonymous with the lack of language influence, as other important factors - such as the newsworthiness of headlines, description of the topic and impact, exaggerations - were not quantified.

In addition to correlations, the high-quality set of PRs was characterized by general features. On average, a press release had a length of 714 words (CI = 681-747). Almost all of them quoted scientists (97.9%, CI = 96.2-99.6). The mean jargon score of PRs was 88.2 (CI = 87.8-88.6), which was on average 10.1 (CI = 9.5-10.7) points higher than the jargon score of matched scientific abstract.

### Engagement of the public in social media

In accordance with the public engagement model, all investigated publications were compared for their popularity on Twitter. Correlation between coverage in news outlets and the number of tweets was moderate (Pearson’s r = 0.571, p<.00001, 95% CI 0.534-0.606). Unsurprisingly, the difference between press-released and not press-released group was large, with a mean of 245 (CI = 186-304) tweets for promoted publications and only 23 (CI = 18-28) tweets for not promoted publications (p<.00001, Supplementary Fig 7).

### Case of CRISPR groundcherries communication

In October 2018, orphan plant Physalis pruinosa was modified with CRISPR to improve productivity traits (Lemmon et al., 2018). This advancement was described in 66 news outlets. It represents the typical pathway of press-release-assisted science communication, where a press release (here: two press releases) about the work was the first non-professional description of the publication on the internet. On the same day as of press release posting, 17 news outlets covered the subject. Coverage continued in 43 news outlets for the next five consecutive days. Over the next two months, there were 6 more news articles.

Both press releases were created by research institutes and contained thorough descriptions with different quotes. Howard Hughes Medical Institute’s press release (1) framed the work as market-centered, focusing on the plant and the possibility of selling groundcherries. It also mentioned intellectual property rights, despite no competing interests declared in the publication. Boyce Thompson Institute’s press release (2) framed the work as more technological, placing emphasis on CRISPR and detailing introduced modifications.

Due to the widespread copying and sublicensing of content, there were 16 original articles. Those articles were compared against each other in a matrix of lexical similarity, which allowed reconstruction of similarity trails (Fig 4). Analysis showed that news articles largely ignored (1) press release. At least 6/16 articles were based in more than half of words on the (2) press release. However, it accounted only for word-for-word similarity and manual inspection revealed that other articles were also influenced. For instance, the 14th article had 40.5% similar words with (2) press release, but the author clearly rewrote multiple sentences from the press release, e.g.:

**Fig 4:**
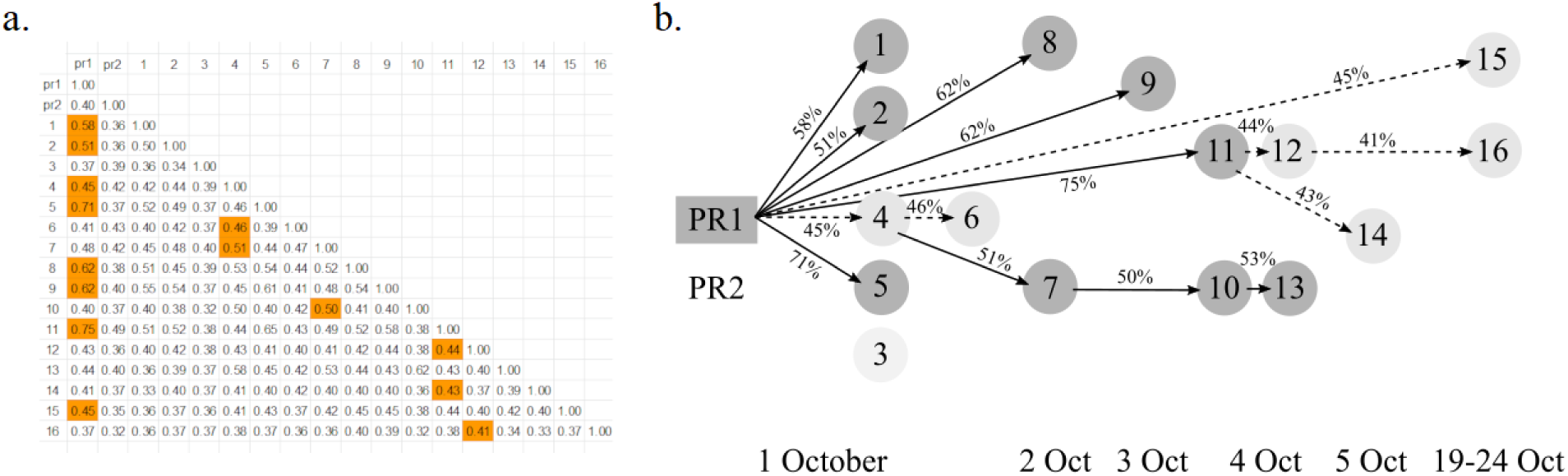
a. Matrix of lexical similarity between all original materials about groundcherries. b. Trails of highest similarity reconstruct probable sources of 16 articles and demonstrate influence of the press release on media coverage.

PR: *Each spring, male deer sprout a new pair of antlers, which are essentially temporary external bones, at a speed unparalleled by the bone growth of other mammals*.

Article: *Every spring, male deer undertake a unique biological ritual: sprouting and rapidly regrowing their massive, spiky antlers*.

This case demonstrated the direct influence of press releases on science communication in online media. 1/3 of original articles were highly similar on a word-for-word basis and some of the remaining news articles concealed copying of the press release with synonyms.

### Case of prime editing communication

In October 2019, prime editing was presented as a novel CRISPR-related technology, allowing a wider range of more precise modifications in the genome (Anzalone et al., 2019). Its intense promotion achieved the goal of world-wide popularity - the finding was described by 166 news outlets, which was surpassed only by 9/1362 of investigated publications - and provided an interesting example of science communication via online media. It is also a valuable account of coverage of competing interests, as the authors applied for a patent on the technology and set up a company intended to profit from therapeutic uses of prime editing.

First public information about prime editing was made available on 11 Oct 2019 at a conference, which was briefly covered in social media (The CRISPR Journal at Twitter, 2019). In the next days, journalists received press briefing under journalistic embargo (agreed moment of publication). It was set for 4:00 PM UTC on 21 Oct 2019. Between 4:02 and 4:10 eight different news outlets published lengthy, original news articles communicating the discovery. Press release was published later, at 4:57, showing that it was a part of the process, but not necessarily the most significant tool. The press release contained disclosure about conflicts of interests, although it did not explicitly mention the application for patent, describing instead “licenses for the use of prime editing”. Following this attitude, only 1/8 early news articles mentioned pending patent application, despite preparing them during the embargo and, for instance, having time to reach and quote independent experts in 6/8 cases. Four more mentioned licenses and company-related conflicts of interest. One article summarized this matter with words: “Liu [senior author] didn’t reply to questions about whether the powerful new tool has any downsides or how financial incentives are shaping the choice to create, and widely share, these means of changing the molecule all life is based on.” (Regalado, 2019)

Prime editing communication was driven by two major points, which were often present in headlines. News articles often framed prime editing as: (1) a cure for the majority of genetic diseases and (2) a tool better than CRISPR. Within embargoed articles, 3/8 based their headlines on (1) and 3/8 based titles on (2). Some pushed those points beyond scientific accuracy, as one article conflated genetic variants with genetic diseases and named the article “A New Crispr Technique Could Fix Almost All Genetic Diseases” (Molteni, 2019). There was also an evident confusion about relation between prime editing and CRISPR - 4 articles described it as “a new CRISPR tool” whereas 4 articles described it as “a tool better than CRISPR”.

This case demonstrated a limited impact of a press release on reporting. Headlines sometimes even contradicted press release’s headlines (point (2)). Lexical similarity between the press release and all initial articles was under half of words despite specialistic vocabulary (42% common words on average). Even if the press release was circulated under embargo, other tools such as press briefings and individual contact had a major influence on the communication of science. Here, the presence of press release is more of proof for engagement of a marketing team, rather than a direct driver of science communication. At the same time, the content of press release was still relevant - about patent application, for instance - because it was written by people who governed the process and reflected their other actions.

### Case of He Jiankui’s experiment communication

In 2018, He Jiankui used CRISPR to modify embryos which were then conceived and became the first genetically modified humans (Cyranoski and Ledford, 2018). Although the story was widely described in virtually all media and in association with multiple sources, the first day of reporting presents a concise and interesting case for sources of science communication. News about this experiment broke on 26 November 2018. Coverage of the study was initiated by the original reporting of MIT Technology Review published at 12:13 AM UTC. Then at 2:48 AM, the second original report was posted by Associated Press. According to accounts of authors (Rathi, 2018), the first report was based on an online investigation, whereas the second report was prepared in advance under embargo agreement with He Jiankui. In addition, He Jiankui prepared six videos, which were made available at 3:41 AM.

Immediately after those reports, 7 major news outlets followed with their coverage. Lexical similarity analysis showed that the second report was the primary source for initial online media coverage, despite later publication. 4/7 early news articles were over 85% word-for-word similar to the second report. Remaining 3/7, which were also chronologically later (4:50 AM, 6:15 AM, 6:52 AM), referred explicitly to both reports, but all of them were more similar to the second report than to the first report. He Jiankui’s videos were quoted in 2/7 early news articles. Later on the same day, at least 34 news outlets published original articles about the matter. Preference of primary sources was preserved: 28/34 (82%) news articles referenced Associated Press as the source and 12/34 (35%) referenced MIT Technology Review. Consistently with explicit references, 30/34 news articles were more similar to the second report, including 7/34 which had more than 50% words in common. Videos of He Jiankui had a substantial impact on coverage on the first day, with 25/34 (73.5%) news articles quoting its parts and often embedding it.

This case demonstrated that original reporting can be prepared and then used similarly as press releases. The first report was largely ignored (not referenced by 2/3 of media), whereas the second report - prepared in advance, with multiple quotes, resembling press release - was the basis of wide coverage in online media. Moreover, videos with the author’s statements were popular among news outlets, essentially serving as the second counterpart of a classic press release.

## Discussion

Assessment of 1362 publications and 461 matched press releases (PRs) showed that the presence of a press release alone determines whether the science is covered in media or not. Publications not promoted through PRs were extremely rarely communicated to the public. Only 7.5% of news articles in popular online news outlets in the US described scientific publications without press releases - lower than in the 1990s in offline press (16% in de Semir, 1998). Even if scientific findings were originally investigated by journalists from renowned magazines (in the present sample: Science, Nature, BBC, New York Times, Washington Post, The Independent), other news outlets did not follow the coverage in absence of PRs. This effect was confirmed in a dataset with high-quality publications - from leading universities and leading journals - and in the unfiltered sample of all research publications related to popular in media CRISPR technology. In the latter, 8.3% of not press-released publications were mentioned on at least one website, but just 1% was covered by general news outlets.

In contrast, press-released science was widely covered in online media. Publications from leading universities or journals were described in 8.8x more news outlets if they had PRs. The unfiltered sample had a more pronounced effect, where the presence of press release led to 74.7x more coverage in online media. Consistently with the large role of PRs in scientific news, their content was used word-for-word in substantial part (39%-43%) of news articles in the most popular online news outlets. This estimate is more conservative - probably due to rewriting and editing - than in the studies of exaggerations in PRs, where inaccurate advice from press release made to 49-58% of associated news articles (Sumner et al., 2014; Bratton et al., 2019).

Worryingly low fraction of press releases disclosed existing competing interests - only 15.7%. Even if the work promoted by PRs was a subject of a patent application, only 1/4 of PRs mentioned this significant conflict of interest. This observation follows the previous study of industry funding disclosures (Woloshin and Schwartz, 2002), which found that 22% PRs acknowledged competing interests, although it was limited to 23 PRs, whereas here 127 PRs were analyzed (including 71 related to intellectual property). As the case of prime editing illustrated, presence and wording of disclosure influences subsequent coverage.

It must be noted that the present investigation is predominantly limited by region (United States), language (English), and topic (genetics and CRISPR). Although geography and language were chosen because of the involvement of US media and universities in the topic, the findings should be explored also in different contexts. The investigated area of science is probably different in comparison to other scientific fields popular in general media, such as artificial intelligence research or psychology. Nonetheless, a wide scope of included scientific findings - from basic to applied - should be representative for the majority of scientific news.

This study established correlational and causal dependence of online news outlets on scientific press releases. The essential question of subject choice in media is almost entirely answered by the presence of press releases - in short, journalists describe publications which are promoted by public information offices. It echoes previous critique of science journalism (Nelkin, 1995; Williams and Gajevic, 2013; Zhang, 2018). The findings pose serious concerns about modern communication of science via media. As the public engagement model assumes communication “about science, including its perils and pitfalls”, press releases are not only - obviously - avoiding shortcomings of research but also rarely acknowledge even the most serious conflicts of interests. In areas of research such as genetics or CRISPR, which are rich in competing interests and bioethical uncertainties, this source of science communication can cause serious consequences, such as overselling science or losing credibility in the eyes of the public.

## Materials and Methods

### Study design

Initial exploratory work assumed another hypothesis: scientifically incorrect headlines of press releases lead to scientifically incorrect coverage in media. Press releases from leading universities were checked for scientific accuracy in headlines. Thorough analysis found a high level of accuracy (84.7% of PRs) and a lack of difference in media coverage between the accurate and incorrect group. This effect was explored further to find out how headlines of PRs translate to headlines of news articles. After collection of the top-down dataset from popular news outlets, similarity analysis of headlines gave an inconclusive result. To ultimately verify the initial line of work, similarity analysis was extended to content. It pointed to a lack of correlation between the use of PRs’ headlines and the use of PRs’ content, which led to the rejection of the first hypothesis. A new hypothesis was formed after additional data about news articles and their subjects were collected (as described in the Results section): not press-released publications are rarely covered in media. This hypothesis was tested in one partially new dataset and one fully new dataset. For the former, previously collected university press releases were used, with the addition of not press-released counterparts, and collection of publications from leading journals, along with data tested in the study and two data points which were not investigated further (use of CRISPR in publications and use of CRISPR keywords in PRs). For the latter, an unbiased set of publications was collected, along with data tested in the study only. Out of three described cases, two were chosen before the study began and one - groundcherries - after analysis of a chain of similarity was performed and showed interesting trails. There was an attempt to extend chains of similarity to all news articles, but it was rejected after statistical objections and inconclusive results in 13/16 checked cases.

### Sample collection

For the top-down dataset (news articles), 14 most popular news outlets were determined on a basis of search engine visibility in Ahrefs tool published in June 2019 (Hardwick, 2019). Although it is not a total measure of popularity (in addition to search engines, people access websites directly and through referrals), search engines are associated with 93% of the internet traffic (Egria and Bayrak, 2014). The list of websites included: nytimes.com, cnn.com, forbes.com, foxnews.com, usatoday.com, businessinsider.com, usnews.com, medicalnewstoday.com, washingtonpost.com, cnet.com, theguardian.com, bbc.com, npr.org. They were searched with Google, Bing, DuckDuckGo with limitation to period 1 Oct 2018 - 30 Sep 2019 for the following keywords in titles: gene, genes, genetics, DNA, genome, genomes, inherited, inheritance, CRISPR. Articles not associated with science (example: financial inheritance) and explicitly declared as opinions were excluded. All other articles were analyzed whether they described a scientific finding: direct link to a publication, mentioned publication, mentioned authors of research, described a study near the date of matched publication.

For the mixed dataset (high-quality publications), 5 leading journals were determined on a basis of impact factors in biomedical sciences from 2018 (New England Journal of Medicine, Lancet, JAMA, Nature, Science), and 50 leading universities were determined on a basis of Times Higher Education ranking labeled as 2020 (Times Higher Education, 2019). Journals and universities were searched with the same keywords as in the previous dataset. Journals were searched with PubMed and Google, universities were searched with Google, Bing, DuckDuckGo with limitation to period 1 Oct 2018 - 30 Sep 2019. Publications labeled as reviews or opinions were excluded, as they do not communicate new scientific findings. Publications without press releases from leading universities were found in the period between 1 Jan 2017 and 30 Sep 2019, searched by senior authors of studies promoted by those universities, expecting that the same scientific group in three years period should have at least one study of sufficiently high quality and newsworthiness. Out of all studies collected for each group, two with the highest popularity (by news outlets and by tweets) from each group were included in the dataset. Here, reviews or opinions were also ignored. Publications common with the previous dataset were excluded.

For the bottom-up dataset (CRISPR publications), all publications containing keyword “CRISPR” in title or abstract were collected with PubMed and Google Scholar from period 1 Oct 2019 - 30 Nov 2019. Publications labeled as reviews or opinions were excluded, as they do not communicate new scientific findings.

### Data collection

The presence of press release was determined by a thorough search using multiple tools (Altmetric, EurekAlert, Google, Ahrefs). Materials labeled as a press release or created by involved parties (university, research organization, publisher, associated non-governmental organization) were included. The number of news outlets covering the publication was determined using Altmetric.com and PlumX in line with Ortega (2018). In several cases PlumX had lower coverage or no information about a publication, therefore instead of averaging both results, the higher number from both measurements was always used. Due to the inconsistent definition of blogs and news outlets in both tools, the number of blogs was always added to the number of news outlets.

Lexical similarity analysis was done using Tool for the Automatic Analysis of Cohesion 2.0.4 (TAACO, Crossley et al., 2016). Although it is a tool designed for inner-text cohesion analysis, it allows an analysis of the similarity between two texts placed as two paragraphs. TAACO compares bags of unique lemmatized words, which is more objective than the direct comparison of text strings. For preprocessing, texts were cleared from interruptions (such as subscription modals, ads, descriptions of images), double new-line signs were replaced with spaces to compress whole text in one line, then compared texts were placed in two paragraphs divided by empty line. The comparison was performed in both directions (text A - text B and text B - text A) and the resulting similarity was a mean of both measurements. The choice of lexical similarity over other methods of comparison (word2vec, LDA) was preceded by calibration experiments with three external (not used in the study) texts describing the same scientific topic.

Acknowledgment of competing interests in press releases was verified by looking for the following keywords: conflict, interest, patent, financial, competing, disclosure. All occurrences were examined manually to confirm if they are referring to conflicts of interest.

Press releases were controlled for promotional keywords from Maat (2007), that is: brand new, top-class, important, large, strong, extensive, terrific, good, special, leading, unique, excellent, reliable, clear, efficient, practical, all, various, several, millions, many, extra, entire, complete, tremendously, considerably, well, strongly, more and more, already, once again, always, constantly, internationally, throughout the world, almost, more than, less than, of course, simply. Additionally, three novelty-specific keywords were looked for: first, novel, innovative. All occurrences were examined manually to ensure their promotional context - phrases such as “first author” or “leading author” were not included.

### Cases

News coverage for all cases was collected from Altmetric.com, PlumX, and Google searches. Exact hour and minute of article posting was determined by searching for the first link to the article at Twitter.com.

### Statistical analysis

All statistical calculations were performed using Jamovi 1.0.7.0 and 1.18.0 (Jamovi Project, 2019; R Core Team, 2018). Files with raw data are available upon request. Confidence levels for all confidence intervals were chosen to be 0.95. All comparisons between groups were performed using Welch’s t-test, as sample sizes were generally highly different and data on news outlets was distributed normally (Shapiro-Wilk p<.00001).

## End Matter

## Acknowledgments

This document was created using an adapted Word preprint template developed by the Finkelstein lab (Finkelstein, 2018).

## Supplementary Materials

### List of articles analyzed in the groundcherries case

https://www.eurekalert.org/pub_releases/2018-10/hhmi-twp092718.php

https://www.eurekalert.org/pub_releases/2018-10/bti-ctt100118.php

https://www.irishexaminer.com/breakingnews/world/groundcherries-at-wimbledon-oh-i-say-872751.html

https://uk.news.yahoo.com/groundcherry-genetically-engineered-fruit-could-150027823.html

https://www.sciencenews.org/article/gene-editing-plant-domestication

https://edition.cnn.com/2018/10/01/health/groundcherry-berry-study/index.html

http://www.xinhuanet.com/english/2018-10/02/c_129964787.htm

https://www.inverse.com/article/49463-how-long-until-i-can-buy-a-pint-of-ground-cherries

https://www.sciencealert.com/meet-weird-fruit-could-soon-become-common-strawberries-future-food-groundcherry-physalispruinosa-crispr

https://www.valuewalk.com/2018/10/groundcherries-marble-sized-fruit/

https://www.techtimes.com/articles/234806/20181003/scientists-want-to-use-gene-editing-to-turn-wild-groundcherry-into-the-nextstrawberry.htm

https://www.usatoday.com/story/news/nation-now/2018/10/03/groundcherry-fruit-aided-crispr-could-hit-grocery-stores/1508764002/

http://www.sci-news.com/biology/groundcherry-next-big-berry-crop-06470.html

https://www.cnbc.com/2018/10/04/tiny-rare-fruit-that-tastes-like-pineapple-could-hit-stores-thanks-to-gene-editing.html

https://weather.com/news/news/2018-10-04-rare-groundcherry-could-reach-grocery-stores

https://www.nytimes.com/2018/10/05/science/groundcherries-crispr-gene-editing.html

https://science.howstuffworks.com/life/botany/rare-groundcherry-gene-editing.htm

https://theconversation.com/tweaking-just-a-few-genes-in-wild-plants-can-create-new-food-crops-but-lets-get-the-regulation-right-104490

### List of articles analyzed in the prime editing case

https://www.bbc.com/news/health-50125843

https://www.statnews.com/2019/10/21/new-crispr-tool-has-potential-to-correct-most-disease-causing-dna-glitches/

https://www.cnet.com/news/breakthrough-gene-editing-tool-can-find-and-replace-dna-better-than-crispr/

https://www.genengnews.com/insights/genome-editing-heads-to-primetime/

https://www.wired.com/story/a-new-crispr-technique-could-fix-many-more-genetic-diseases/

https://www.newscientist.com/article/2220476-crispr-upgrade-could-make-genome-editing-better-and-safer/

https://www.technologyreview.com/s/614599/the-newest-gene-editor-radically-improves-on-crispr/uses

https://www.sciencemag.org/news/2019/10/new-prime-genome-editor-could-surpass-crispr

https://www.broadinstitute.org/news/new-crispr-genome-editing-system-offers-wide-range-versatility-human-cells

### List of articles analyzed in the He Jiankui case

https://www.technologyreview.com/s/612458/exclusive-chinese-scientists-are-creating-crispr-babies/

https://apnews.com/4997bb7aa36c45449b488e19ac83e86d

https://www.foxnews.com/health/chinese-researcher-claims-to-have-altered-babies-dna

https://www.dailymail.co.uk/news/article-6428275/First-gene-edited-babies-claimed-China.html

https://www.bloomberg.com/news/articles/2018-11-26/urgent-ap-exclusive-first-gene-edited-babies-claimed-in-china

https://www.statnews.com/2018/11/25/china-first-gene-edited-babies-born/

https://techcrunch.com/2018/11/25/crispr-scientist-in-china-claims-his-teams-research-has-resulted-in-the-worlds-first-gene-editedbabies/

https://www.geekwire.com/2018/chinese-scientist-says-first-gene-edited-babies-born-effort-fight-hiv/

https://www.cnet.com/news/scientists-in-china-claim-to-have-created-first-gene-edited-human-babies/

https://www.youtube.com/watch?v=th0vnOmFltc

https://www.statnews.com/2018/11/26/he-jiankui-gene-edited-babies-china/

https://www.theguardian.com/science/2018/nov/26/worlds-first-gene-edited-babies-created-in-china-claims-scientist

https://www.nature.com/articles/d41586-018-07545-0

https://www.nytimes.com/2018/11/26/health/gene-editing-babies-china.html

https://www.sciencemag.org/news/2018/11/crispr-bombshell-chinese-researcher-claims-have-created-gene-edited-twins

https://www.npr.org/sections/health-shots/2018/11/26/670752865/chinese-scientist-says-hes-first-to-genetically-edit-babies

https://qz.com/1474384/chinas-gene-edited-crispr-babies-push-bioethics-into-a-dark-new-era/

https://www.theatlantic.com/science/archive/2018/11/first-gene-edited-babies-have-allegedly-been-born-in-china/576661/

https://www.nbcnews.com/health/health-news/chinese-scientist-says-he-made-gene-edited-twins-using-crispr-n940026

https://www.bbc.com/news/health-46342195

https://www.newscientist.com/article/2186504-worlds-first-gene-edited-babies-announced-by-a-scientist-in-china/

https://arstechnica.com/science/2018/11/chinese-scientist-claims-to-have-gene-edited-humans/

https://www.reuters.com/article/us-health-china-babies-genes/chinese-university-to-investigate-after-academic-claims-to-haveedited-twins-genes-idUSKCN1NV19T

https://www.washingtonpost.com/science/2018/11/26/scientists-claim-gene-edited-babies-creates-uproar/

https://www.popularmechanics.com/science/health/a25305651/he-jiankui-crispr-genetic-editing-twins-aids-hiv/

https://futurism.com/chinese-researchers-gene-edited-human-embryos-hiv-crispr

https://www.ft.com/content/4bc28f80-f142-11e8-ae55-df4bf40f9d0d

https://theconversation.com/worlds-first-gene-edited-babies-premature-dangerous-and-irresponsible-107642

https://www.the-scientist.com/news-opinion/claim-of-first-gene-edited-babies-triggers-investigation-65139

https://www.france24.com/en/20181126-china-hiv-scientist-claim-gene-edited-babies-dna

https://www.theverge.com/2018/11/26/18112970/crispr-china-babies-embryos-genetic-engineering-bioethics-policy

https://www.independent.co.uk/news/world/asia/china-babies-genetically-edited-altered-twins-scientist-dna-crispr-a8651536.html

https://www.buzzfeednews.com/article/briannasacks/chinese-scientist-gene-edited-babies

https://www.discovermagazine.com/health/chinese-scientist-claims-hes-created-worlds-first-gene-edited-babies

https://www.genomeweb.com/scan/researcher-claims-first-gene-edited-infants

https://phys.org/news/2018-11-gene-edited-baby-chinese-scientist-outrage.html

https://www.livescience.com/64166-first-genetically-modified-babies-risks.html

https://www.sciencealert.com/one-scientist-is-claiming-the-first-babies-from-crispr-edited-embryos-have-already-been-born

https://theconversation.com/researcher-claims-crispr-edited-twins-are-born-how-will-science-respond-107693

https://www.iflscience.com/health-and-medicine/chinese-scientist-claims-to-have-created-first-geneedited-babies/

https://gizmodo.com/reports-of-first-genetically-enhanced-babies-spark-outr-1830657274

https://fortune.com/2018/11/26/chinese-researchers-genetic-engineering-babies-crispr-hiv/

https://www.geek.com/news/chinese-scientist-claims-he-made-worlds-first-genetically-edited-babies-1762741/

https://www.buzzfeednews.com/article/nidhisubbaraman/hiv-crispr-china-twins

## Supplementary Figures

**Supplementary Fig 1:**
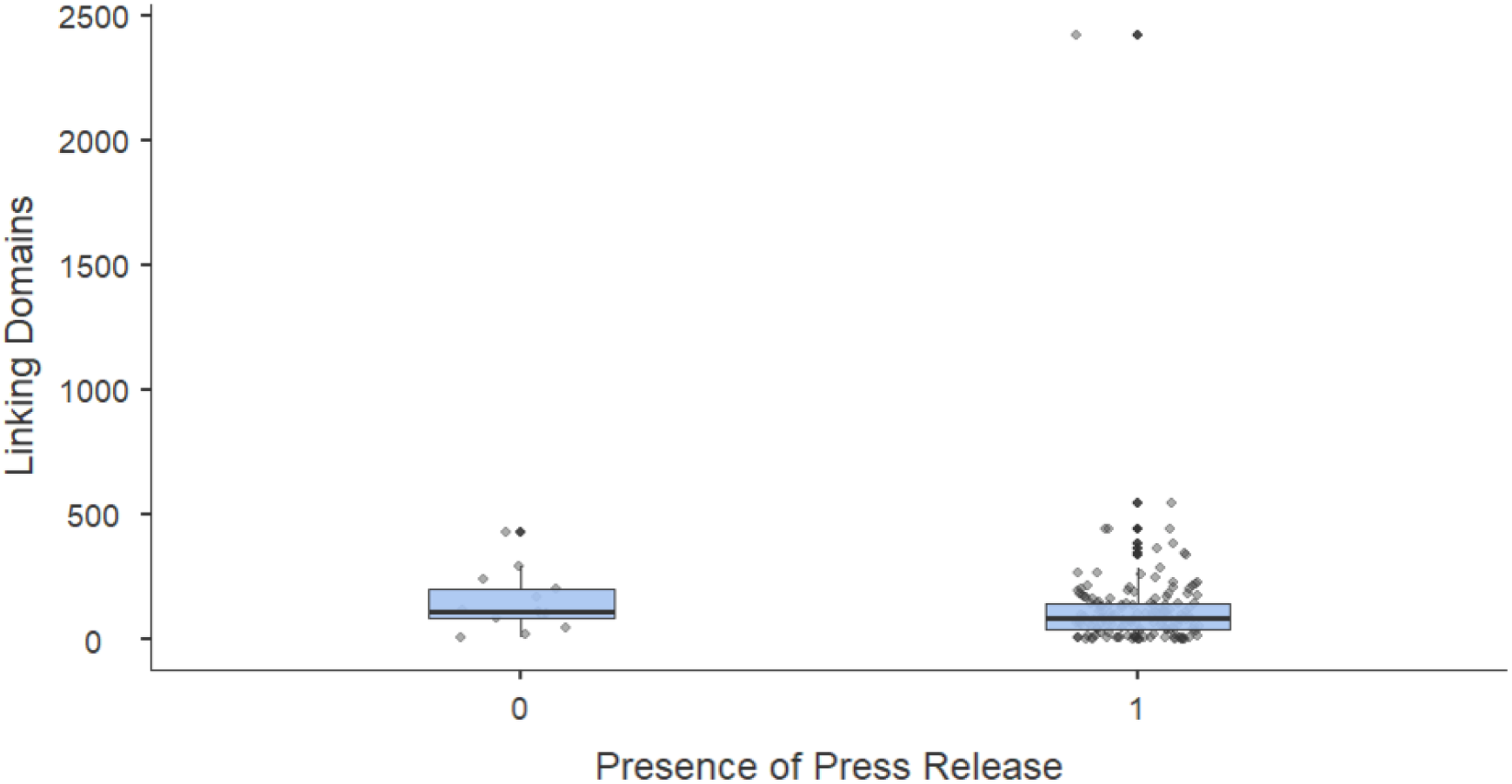
Box plots with all data points of news articles popularity, measured by number of linking domains, divided to group of press-released and not press-released subjects.

**Supplementary Fig 2:**
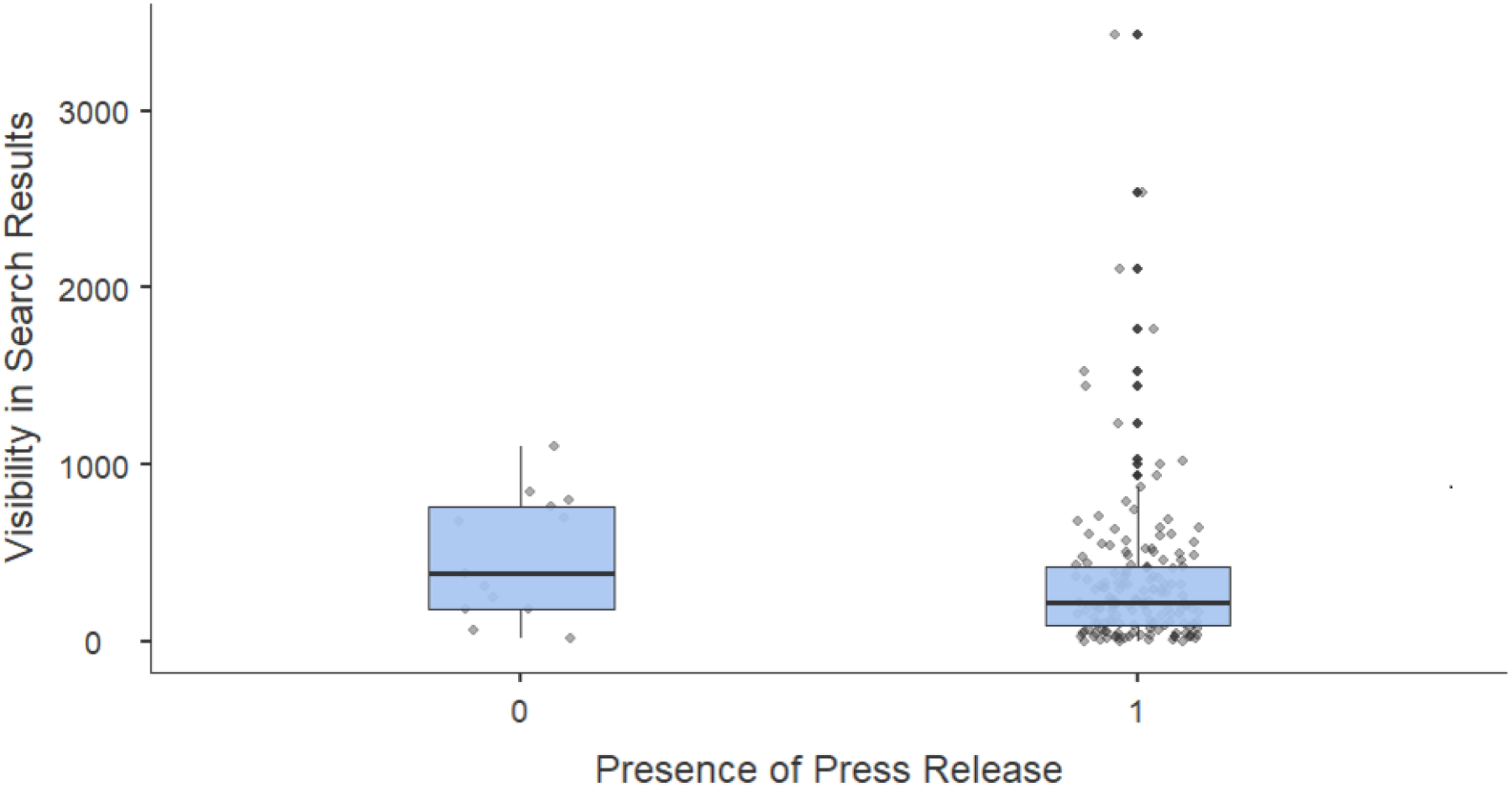
Box plots with all data points of news articles popularity, measured by number of phrases they visible under, divided to group of press-released and not press-released subjects.

**Supplementary Fig 3:**
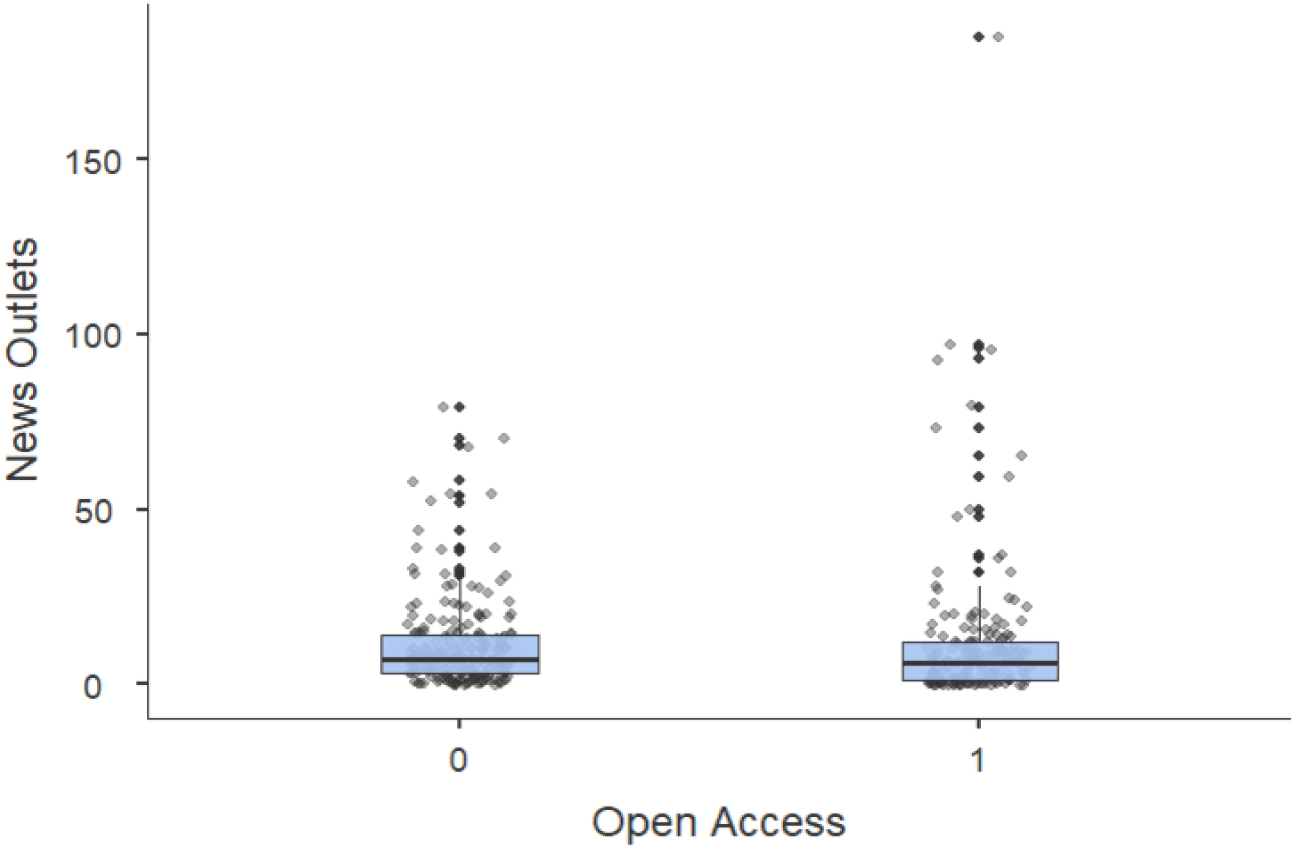
Box plots with all data points on news outlet number, divided by accessibility.

**Supplementary Fig 4:**
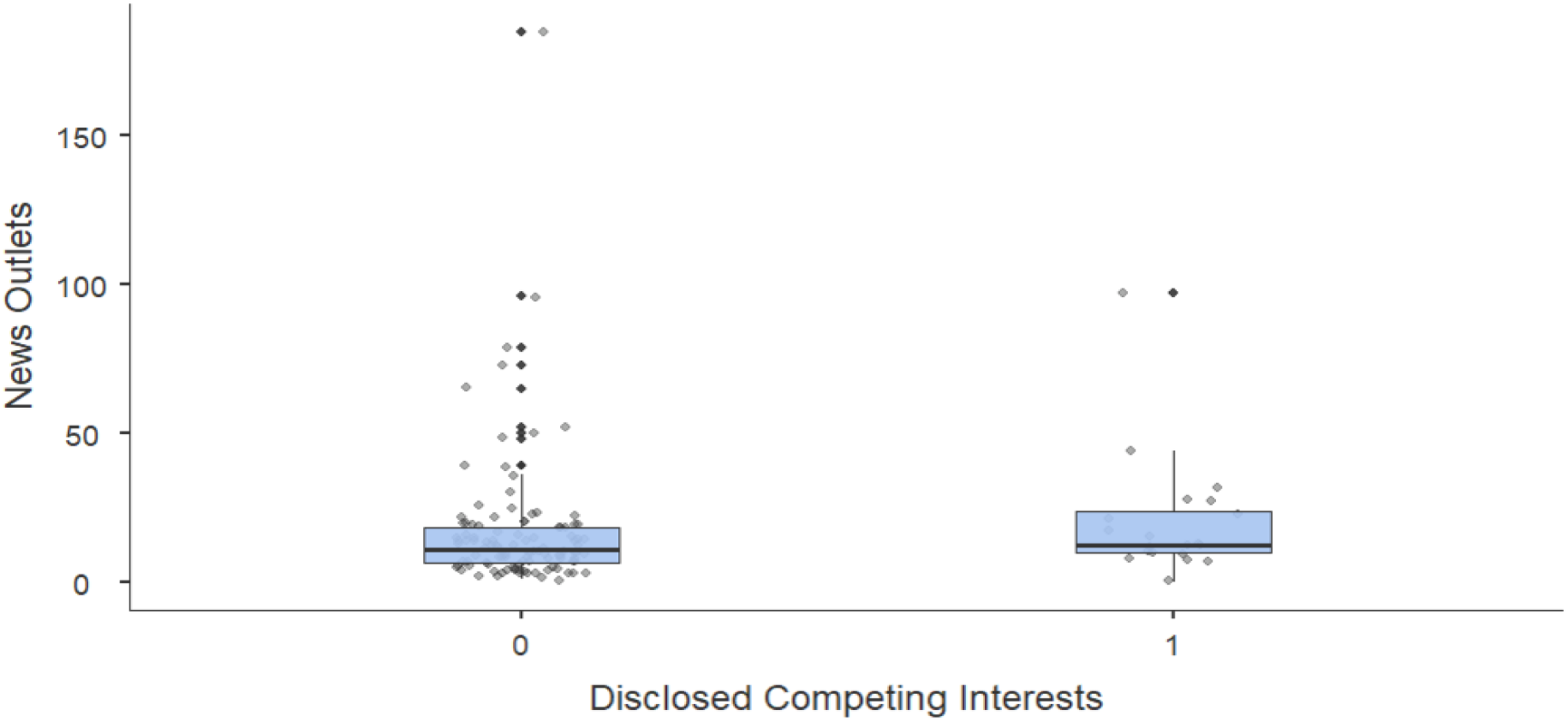
Box plots with all data points of news outlet number, divided by disclosure of competing interests.

**Supplementary Fig 5:**
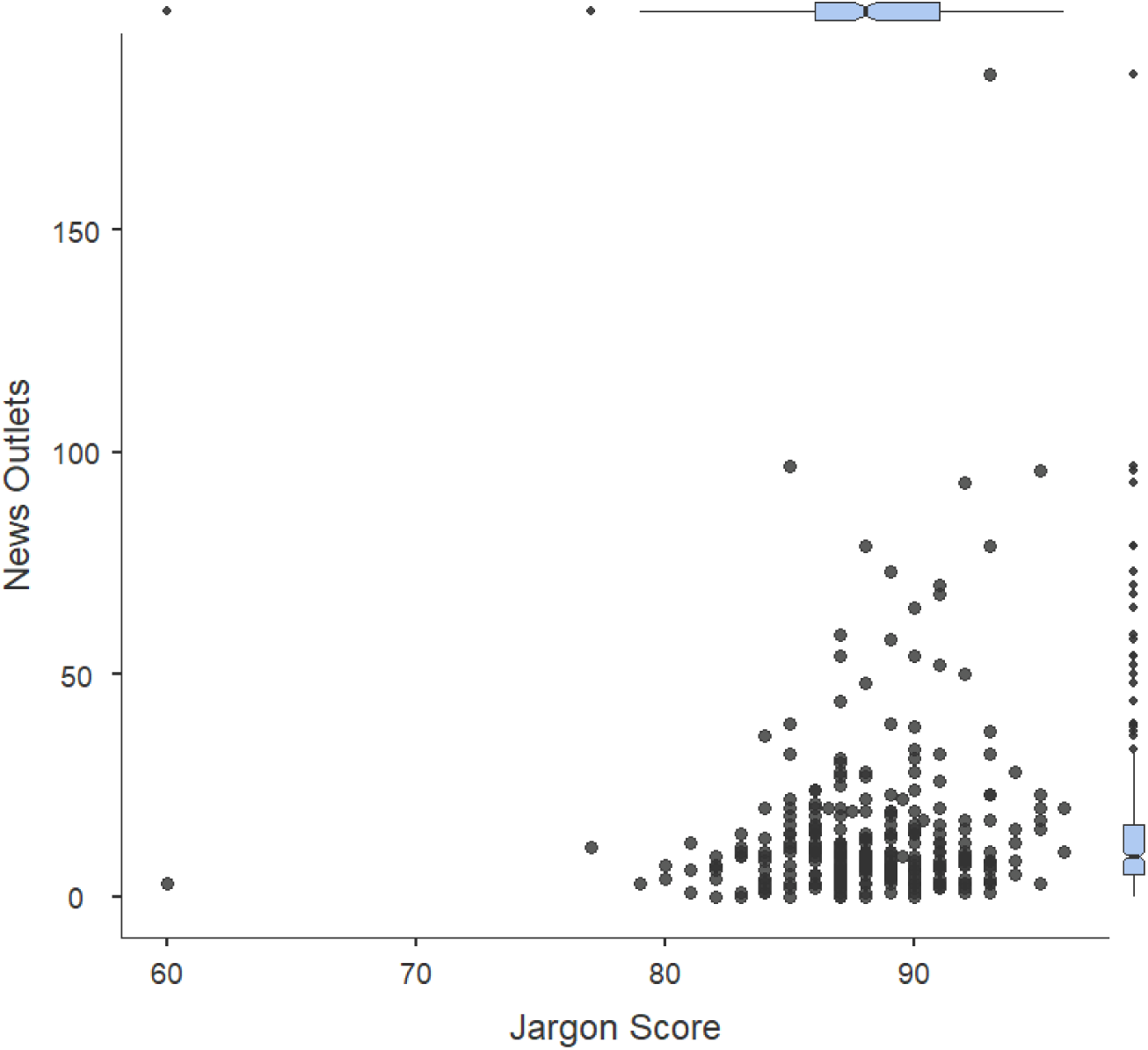
Scatter plot of news popularity and jargon scores of source press releases.

**Supplementary Fig 6:**
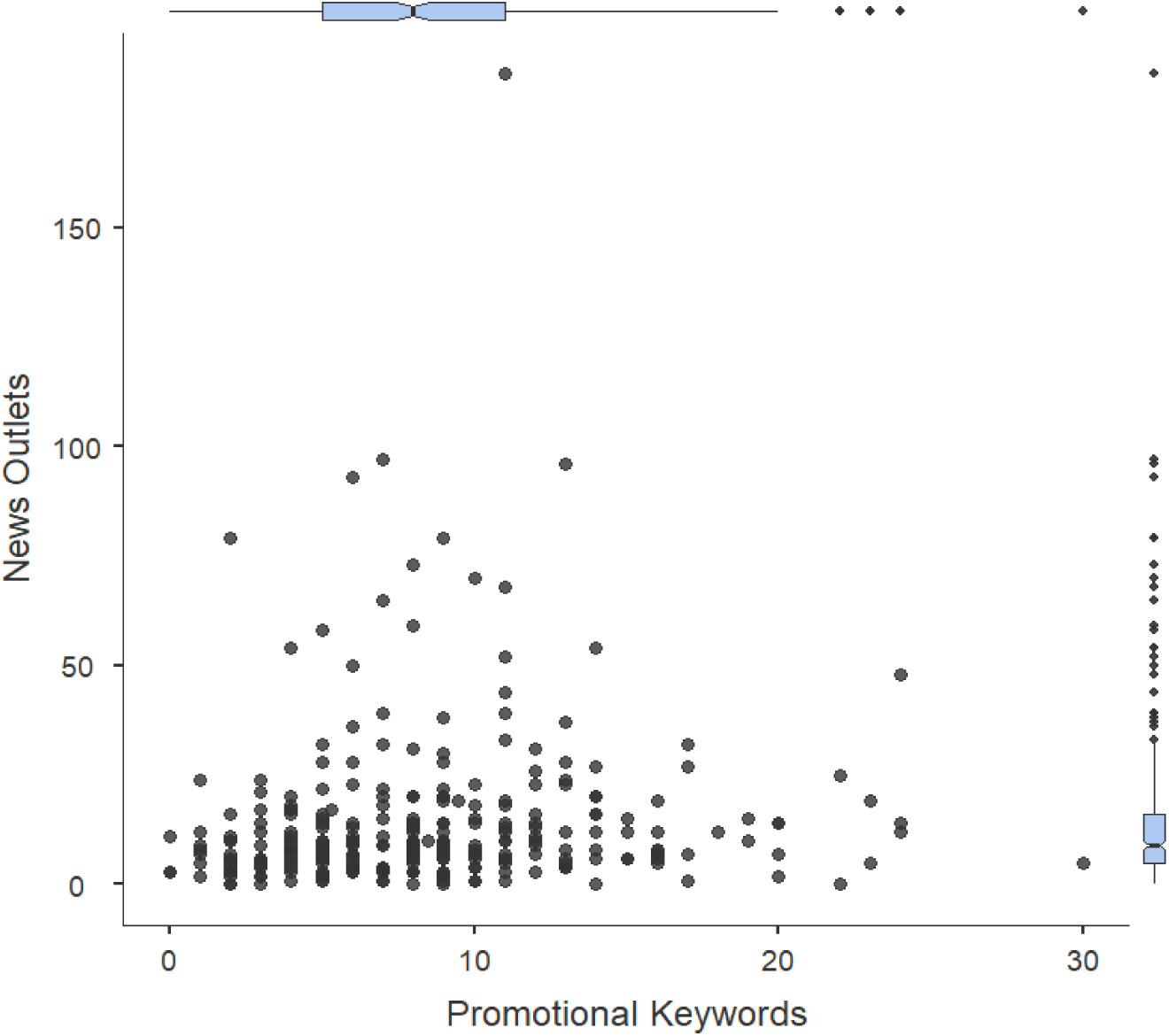
Scatter plot of news popularity and number of promotional keywords in source press releases.

**Supplementary Fig 7:**
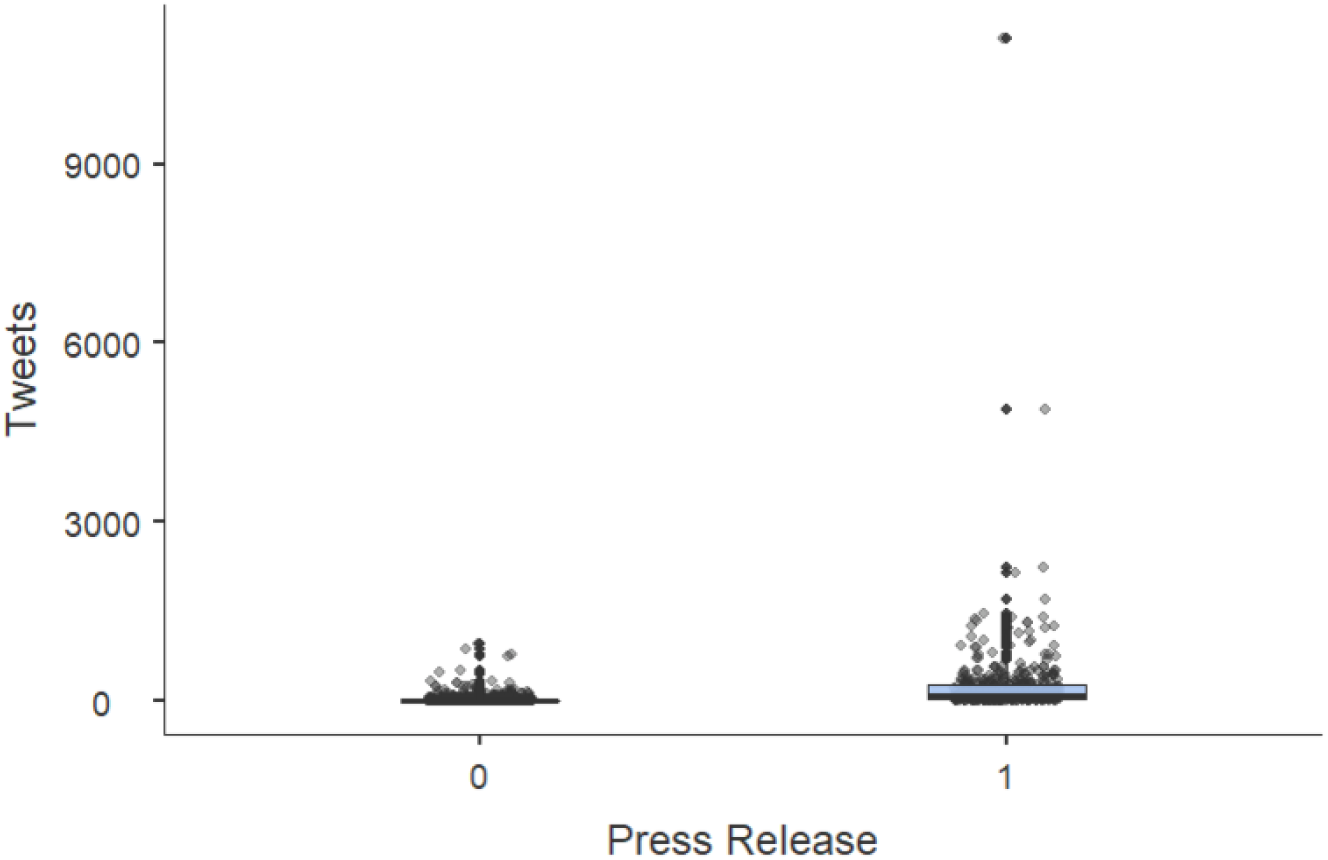
Comparison of social media engagement between press-released and not press-released subjects.

